# Striatal iron content is linked to reduced fronto-striatal brain function under working memory load

**DOI:** 10.1101/812602

**Authors:** Karen M. Rodrigue, Ana M. Daugherty, Chris M. Foster, Kristen M. Kennedy

## Abstract

Non-heme iron accumulation contributes to age-related decline in brain structure and cognition via a cascade of oxidative stress and inflammation, although its effect on brain function is largely unexplored. Thus, we examine the impact of striatal iron on dynamic range of BOLD modulation to working memory load. N=166 healthy adults (age 20-94) underwent cognitive testing and an imaging session including n-back (0-, 2-, 3-, and 4-back fMRI), R2*-weighted imaging, and pcASL to measure cerebral blood flow. A statistical model was constructed to predict voxelwise BOLD modulation by age, striatal iron content and an age × iron interaction, controlling for cerebral blood flow, sex, and task response time. A significant interaction between age and striatal iron content on BOLD modulation was found selectively in the putamen, caudate, and inferior frontal gyrus. Greater iron was associated with reduced modulation to difficulty, particularly in middle-aged and younger adults with greater iron content. Further, iron-related decreases in modulation were associated with poorer executive function in an age-dependent manner. These results suggest that iron may contribute to differences in functional brain activation prior to older adulthood, highlighting the potential role of iron as an early factor contributing to trajectories of functional brain aging.

## 1. Introduction

Iron is the most prevalent metal in the brain and numerous postmortem and *in vivo* studies have demonstrated that in older adult brains, concentration of metabolically-active iron particles is increased in cortical and subcortical gray matter (e.g., Hallgren & Sourander, 1958; Haacke et al., 2005; Bartzokis et al., 2007). Iron is a necessary metabolite for normal cell functions, including adenosine triphosphate (ATP) production, DNA replication, and in the brain, myelination, neurotransmitter production and expression (Zecca et al. 2004; Ward et al. 2014). Proper regulation of iron homeostasis is critical to healthy brain function, and non-heme iron accumulation outside of regulatory proteins promotes free radical formation, which in excess constitutes oxidative stress (Winterbourn 1995). In this manner, excess accumulation of non-heme iron precipitates mitochondrial dysfunction, induces inflammation, and accelerates apoptosis (Zecca et al. 2004).

Iron concentration can be measured non-invasively from *in vivo* MRI, and an expanding literature clearly demonstrates an age-related increase in regional brain iron in even healthy aging (Rodrigue et al. 2011; Daugherty and Raz 2013, 2015). Further, elevated iron concentration predicts regional brain atrophy (Daugherty et al. 2015; Daugherty and Raz 2016), and deficits in behavioral and cognitive performance that are typical in aging (Sullivan et al. 2009; Rodrigue et al. 2013; Daugherty and Raz 2015; Daugherty et al. 2019). Therefore, iron accumulation appears to be an antecedent to metabolic dysfunction and inflammation that drives progressive declines in brain structure and cognition in aging (Raz and Daugherty 2018).

However, relatively little is known about the effects of iron accumulation on brain function. To date, there is a single report linking iron concentration and task-based fMRI blood oxygen-level dependent (BOLD) signal (Kalpouzos et al. 2017) and another report of iron concentration correlating with task-free functional coherence (Salami et al. 2018). These recent fMRI findings demonstrate that higher striatal iron content is associated with both decreases in fronto-striatal task-based activation during a mental imagery task (Kalpouzos et al. 2017), in addition to lower spontaneous coherence within caudate nucleus and putamen resting state networks and decreases in the number of functional connections between the putamen and other networks (Salami et al. 2018).

The striatum are among the most iron-rich regions in the brain (Hallgren and Sourander 1958) and preferential accumulation of iron in the striatum across the lifespan predicts volumetric atrophy and cognitive decline years later (Daugherty et al. 2015; Daugherty and Raz 2016, 2017). Striatal nuclei support cognitive performance via fronto-striatal loops (Alexander et al. 1986). Executive functions (EF) and working memory decline with age likely due to the relative vulnerability of both the striatum and the dorsolateral prefrontal cortex (PFC) to the aging process (Tekin and Cummings 2002). Therefore, it is plausible that iron accumulation within the striatum throughout the lifespan may contribute to changes in fronto-striatal function to impact executive function and working memory performance.

The preliminary evidence of functional brain activation differences during task and at rest (Kalpouzos et al. 2017; Salami et al. 2018) offer further support of brain iron accumulation as a risk factor for age-related neural and cognitive decline. Given the limited research thus far, it is unclear when in the adult lifespan striatal brain iron accumulation may begin to contribute to the functional activation changes observed in aging. Further, it is unknown if iron content effects broader task-relevant processes (i.e., modulation of activation to increased cognitive challenge; Kennedy et al., 2015; Kennedy, Boylan, Rieck, Foster, & Rodrigue, 2017; Rieck, Rodrigue, Boylan, & Kennedy, 2017), and how these together relate to cognitive performance across the lifespan.

Here, we estimate regional iron content *in vivo* from high-resolution R2*-weighted magnetic resonance imaging (MRI), and test striatal iron concentration as a predictor of BOLD modulation to increasing cognitive challenge across the adult lifespan (*N* = 166 healthy adults, age 20-94 years). We hypothesized that increased striatal iron concentration would predict poorer modulation of BOLD activation to working memory load, and that this iron-related reduction in BOLD modulation would be associated with poorer cognitive outcome.

## 2. Methods

### 2.1 Participants

Participants included 166 healthy adults (mean age = 52.75 ± 19.06; age range 20 – 94; see **Table 1**), drawn from a larger sample of 180 individuals recruited from the Dallas-Fort Worth metroplex who had complete fMRI and R2* data. Of those 180, a total of 14 participants’ data were excluded from analysis due to: excessive in-scanner motion during fMRI (*n* = 3), poor quality anatomical scans (*n* = 2), poor quality functional scans (*n* = 3), distortion or motion in R2* images (*n* = 4), responding to less than 70% of fMRI trials (*n* = 1), and an extreme value relative to the mean on R2* (> 7 SD; *n* = 1). Participants were screened against a history of metabolic, neurological or psychiatric conditions, head trauma, drug or alcohol problems, significant cardiovascular disease, depression (Center for Epidemiological Study - Depression < 16 (Radloff 1977), and to be cognitively intact (Mini Mental State Exam ≥ 26 Folstein, Folstein, & McHugh, 1975). Thirty-five individuals self-reported a history of hypertension.

**Table 1.** Participant Demographics and Task Performance (Mean ± SD)

|  |  |
| --- | --- |
| N (% Women) | 166 (40.96) |
| Age | 52.75 (19.06) |
| Education | 15.63 (2.49) |
| MMSE | 29.04 (0.86) |
| CESD | 4.14 (3.77) |
| CBF | 23.37 (5.16) |
| Striatal R2* | 28.06 (6.20) |
| fMRI Task Accuracy |  |
| Zero-Back | 94.55 (8.69) |
| Two-Back | 85.31 (11.75) |
| Three-Back | 78.33 (9.78) |
| Four-Back | 76.88 (8.58) |
| fMRI Task RT (sec) |  |
| Zero-Back | 0.65 (0.13) |
| Two-Back | 0.87 (0.22) |
| Three-Back | 0.96 (0.23) |
| Four-Back | 0.96 (0.25) |

*Note:* Both whole brain mean cerebral blood flow (CBF), $r(166) = -.28$ , $p < .001$ , and iron (R2\*), $r(166) = .58$ , $p < .001$ , exhibit significant relationships with age. MMSE = Mini Mental State Exam; CESD = Center for Epidemiological Study – Depression Scale; CBF = cerebral blood flow; RT = Response Time; unless otherwise noted, numbers in parentheses represent standard deviations.

### 2.2 Cognitive Measures

Prior to MRI scanning, participants underwent two days of cognitive testing in which a variety of cognitive measures were collected. The current study focuses on executive function and working memory measures, detailed below.

#### 2.2.1 Executive Function (EF)

The executive function measures consisted of several subtests of the Delis-Kaplan Executive Function System (Delis et al. 2001) including Verbal Fluency (subtests: category fluency – total average correct of animals and boy’s names; verbal fluency – average total correct for letters F, A, S), the Trail Making Test (subtests: trail switching adjusted for average time on trail numbers and trail letters), and the Color Word Interference Test (subtests: color-word inhibition adjusted for average time on color naming and color reading; color-word inhibition switching adjusted for average time on color naming and color reading). Subtests from the Wisconsin Card Sorting Test (WCST; Heaton, Chelune, Talley, Kay, & Curtiss, 1993) (subscores: number of categories completed, number of trials to complete first category, and number of failures to maintain set) were also administered.

#### 2.2.2 Working Memory (WM)

The working memory measures consisted of Wechsler Adult Intelligence Scale – IV (Wechsler 2008) subtests Digit Span forward and backward (subscores: number correct on each test), and the Listening Span task (Salthouse et al. 1991) (subscores: simple span, absolute span, and total span).

To create reliable cognitive constructs for EF and WM, we utilized composite measures of executive function scores and working memory scores, respectively. All tests scores were standardized to *z*-scores before calculating the composites. Cronbach’s α = .71 and .89, for the EF and WM composite, respectively. These composite scores were used to investigate the effects of age, striatal iron content and BOLD modulation on these two domains of cognition.

### 2.3 Neuroimaging Protocol

#### 2.3.1 MRI sequence acquisition

All participants were scanned on a single 3T Philips Achieva scanner equipped with a 32-channel head coil. All images were collected parallel to the AC-PC line. For task-evoked fMRI, Blood Oxygenation Level Dependent (BOLD) data were collected using a T2*-weighted echo-planar imaging (EPI) sequence with 29 interleaved axial slices per volume providing full brain coverage, (64 × 64 × 29 matrix, voxel size = 3.44 × 3.44 × 5 mm^3^, FOV = 220 mm^2^, TE = 30 ms, TR = 1.5 s, flip angle = 60°, 20 min). High resolution anatomical images were acquired using a T1-weighted MP-RAGE sequence (160 sagittal slices, 1 × 1 × 1 mm^3^ voxel size; 256 × 204 × 160 matrix, TR =8.3 ms, TE= 3.8 ms, flip angle = 12°, 3:57 min). For measurement of regional iron content, a T2*weighted multi-echo 3D gradient-recalled echo (GRE) sequence was acquired (65 axial slices, eight echo times: 5.68 ms +Δ 2.57 ms, FA = 15°, TR = 37 ms, FOV = 256 mm^2^, 512 × 512 × 65 matrix, voxel size = 0.5 × 0.5 × 2 mm^3^, 10:14 min). For a measure of cerebral blood flow, arterial spin labeled images were acquired using a pcASL sequence (labeling time = 1516 ms, postlabeling delay = 1500 ms, labeling plane offset = 20 mm, two-dimensional pcASL readout parameters TR = 4460 ms, TE = 17 ms, 64 × 64 × 29 matrix; voxel size = 3.44 × 3.44 × 5 mm^3^, FOV = 220 mm^2^, EPI factor = 35, 35 label-control pairs, M0 image = mean of control images, 5:20 min).

#### 2.3.2 Data Processing for Regional Iron Estimates from R2*

Procedures for processing the T2* multi-echo GRE data for estimates of regional iron content are detailed in prior reports (Rodrigue et al. 2013; Daugherty et al. 2015, 2019). Briefly, images were processed and masked in the Signal Processing in NMR software package (SPIN; MR Innovations, Inc., Detroit, MI, USA: http://mrinnovations.com/spin-lite; last accessed 03/29/2019). R2* was calculated at each voxel as the inverse of T2*, for which higher intensity values indicate relatively greater iron content. A statistical threshold derived from the Gaussian distribution of the observed intensity values across the multi-echo image set was applied during R2* map creation to improve signal-to-noise ratio. Estimates of iron content were sampled from the bilateral caudate nucleus and putamen. We chose to sample from the neostriatum given the relative salience of these regions to higher cognitive function such as working memory versus palleostriatum. Standardized, 24-pixel circular masks were manually placed within the regions of interest (see **Figure 1**). On a single image slice, four masks were placed within a region, per hemisphere. The manual placement of several, smaller masks provided a representative estimate of average iron content throughout the region, while avoiding vascular objects (i.e., heme iron), and partial voluming with white matter or cerebrospinal fluid, all of which would bias the estimate. High iron concentrations within the striatum bias regional segmentation (Lorio et al. 2014) and the use of standardized masks minimized this source of bias. In this manner, each region was measured on three contiguous, 2-mm slices and the bilateral average estimate per region was taken from a total of 24 standardized masks (left and right combined) placed throughout the middle of each region (see Daugherty et al., 2019 for detailed procedure). Reliability was established between two raters in a sample of 10 participants and intra-class correlation coefficients (Shrout and Fleiss 1979) were greater than 0.90 for each region in each hemisphere. In the reported analyses, iron estimates were averaged across the regions of interest to calculate an estimate of striatal iron content.

**Figure 1.**
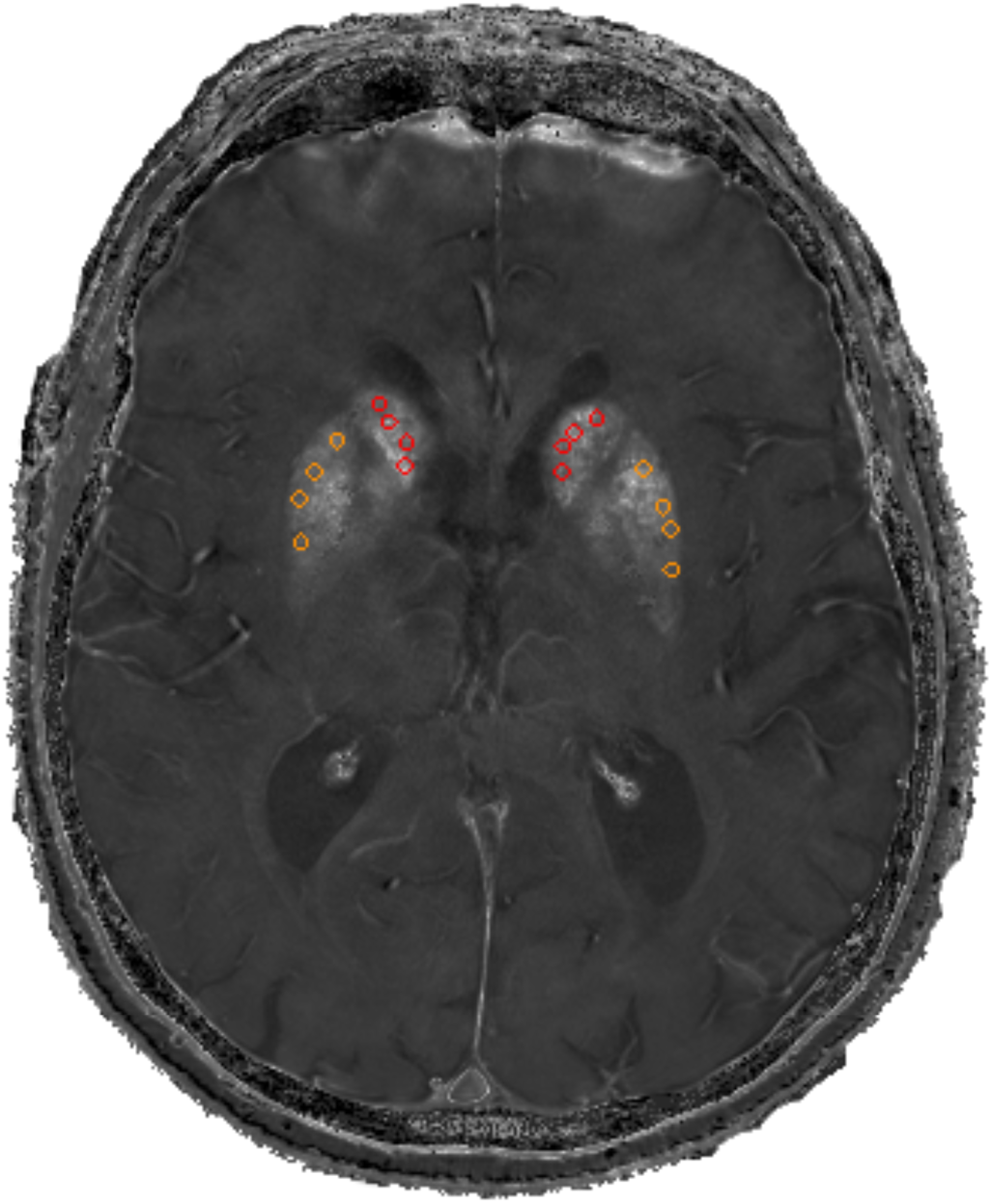
Illustration of masks within the striatum overlaid on an R2* map. A representative axial slice through the basal ganglia from the R2* image showing manually drawn masks in the Caudate (red) and Putamen (orange). Four masks were drawn in each hemisphere of a region and repeated on 3 consecutive axial slices. Voxels within the 12 masks per region, per hemisphere, were averaged to create the index of striatal iron content.

#### 2.3.3 ASL Data Processing

Each individual’s ASL scan was processed as follows. The 70 dynamic volume images were split into separate tag/labeled and control images, with each image containing 35 volumes. A difference image was calculated for each of the labeled and control images, which was then used as input to calculate perfusion images using the Bayesian Inference for Arterial Spin Labeling MRI (BASIL) toolset (Chappell et al. 2009), which is available as part of the FMRIB Software Library (fsl.fmrib.ox.ac.uk/fsl/fslwiki/BASIL). Cerebral blood flow (CBF) images were corrected for partial voluming within BASIL and indicate blood flow in each grey matter voxel (ml/100gm/min). These images were then co-registered to the individual participant’s T1 MP-RAGE image. T1-MPRAGE images were processed using the semi-automated Freesurfer v5.3 pipeline (Fischl and Dale 2000) to generate all grey matter parcels from the Desikan atlas (Desikan et al. 2006) for each individual. Each individual’s Freesurfer parcels were visually inspected and manually edited when necessary to ensure accurate segmentation of grey matter and then warped to individual participant ASL space. Whole brain CBF was computed as the average of all parcels from the partial volume corrected CBF image. This whole brain CBF measure was used as a covariate in the fMRI analysis, to control for the potential mutual contribution of heme iron in circulating blood to both the R2*and BOLD measures.

#### 2.3.4 fMRI N-Back task

Using a block design, participants were presented with an *n*-back task using digits as stimuli. For each trial, participants made a SAME/DIFFERENT response with their index (same) or middle (different) finger to indicate whether the digit was the same or different as *n* digits ago. Before each block began, a 5 sec cue indicated which type of *n*-back was about to start: 0-back, 2-back, 3-back, or 4-back, followed by 2 sec of fixation prior to the presentation of digits. For the 0-back trials, participants made a digit identification decision, indicating whether or not the digit on the screen was the one cued at the beginning of the block. Digits (“2-9”) were presented in six pseudo-counterbalanced blocks, for 500 ms with a 2000 ms inter-stimulus interval using Psychopy v1.77.02 (Peirce 2007, 2009). Each run of data consisted of 8 blocks, including two blocks of each level of difficulty. The blocks were counterbalanced for difficulty within run. Of the 420 trials, 144 were match trials (18 for 0-back and 42 each for 2-,3-,4-back), and 276 were non-match trials (42 0-back and 78 each for 2-,3-,4-back). The 0-back blocks had 10 trials, while the 2-, 3-, and 4-back blocks each had 20 trials. The trials were presented in a pseudo-random order. There were three runs total, yielding a total functional scan time of about 20 minutes.

#### 2.3.5 fMRI Data Processing

Data preprocessing and statistical analyses were performed using SPM8 (Wellcome Department of Cognitive Neurology, London, UK) along with in-house Matlab R2012b (Mathworks) scripts. Additionally, Art Repair toolbox (Mazaika et al. 2007) was used to identify potential outliers in movement (> 2 mm displacement) and intensity shift (> 3% deviation from the mean in global intensity spikes) in the EPI images. Runs with >15% of total volumes (∼40 volumes) marked as outliers for movement were excluded (*n* = 3). In order to be included in the analysis, participants were required to have at least two runs (out of three runs total) with quality data. Functional images were adjusted for slice acquisition time and motion correction using 6 directions of motion-estimates from ArtRepair included as nuisance regressors at the subject level, and each participant’s T1-weighted anatomical image was used to co-register the functional maps to standardized MNI space. The resulting normalized images were smoothed with an isotropic 8mm FWHM Gaussian kernel. These procedures were identical to Kennedy et al., (2017). Beta estimates derived as a mean from all voxels within a cluster were extracted using a cluster mask on each subject’s first-level effects.

### 2.4 Experimental Design and Statistical Analyses

#### 2.4.1 Task Pre-training Procedure

Participants were trained on the task just prior to entering the scanner to maximize their ability to understand task instructions and to complete the four levels of working memory load. First, trained researchers demonstrated the task, including a real-time on-screen run-through of each level of difficulty, as well as a schematic detailing when participants should respond “same” or “different”. Once the participant completed all levels of difficulty s/he completed a second brief practice to emulate what they would experience in the scanner (cue presentation, timing, etc.). The participants had on average 7.13 (± 2.26; range 5-16) exposures during practice. Participants also completed a post-scan questionnaire providing information about their experience in the experiment.

#### 2.4.2 fMRI Data Analysis

Subject-level and group-level analyses were performed using the general linear model in SPM8. At the individual subject level, regressors for each level of difficulty were created (0-, 2-, 3-, 4-back), and a linear contrast was computed across level of *n*-back (using −2.25, −0.25, 0.75, 1.75 weights), reflecting the slope of change in BOLD as a function of increasing working memory load. For the group-level analyses, age, iron, the age × iron interaction, whole brain average CBF, sex, and average task response time were used as between-subjects second-level covariates, predicting linear modulation from the first level model. To alleviate bias in regression coefficients from multicollinearity, all covariates were mean-centered (Irwin and McClelland, 2001) except for sex and the interaction term between age and striatal iron. Two primary contrasts of interest were calculated. First, we examined parametric increases and decreases in activation to increasing WM load (positive modulation effect and negative modulation effect). Second, we examined the age × iron interaction on modulation to difficulty (WM load) to identify regions where modulation depended upon iron and age. A height threshold of *p* < .001 was chosen for the primary analysis and reported cluster *p*-values; however, we additionally present results with a height threshold of *p* < .005 in Figure 2 for illustration of the more generalized effect. To ensure that Type I error rates were correctly controlled at the cluster level (Eklund et al. 2016), cluster corrections were calculated using the Statistical nonParametric Mapping toolbox (SnPM13; http://warwick.ac.uk/snpm) with 5000 permutations to derive the *FWE* corrected clusterwise threshold. Importantly, non-parametric methods, such as the permutation implemented in SnPM13, make no assumption about the underlying spatial auto-correlation function and thus make accurate clusterwise inferences regardless of the chosen cluster-defining threshold. These are the cluster FWEs presented in coordinate tables.

**Figure 2.**
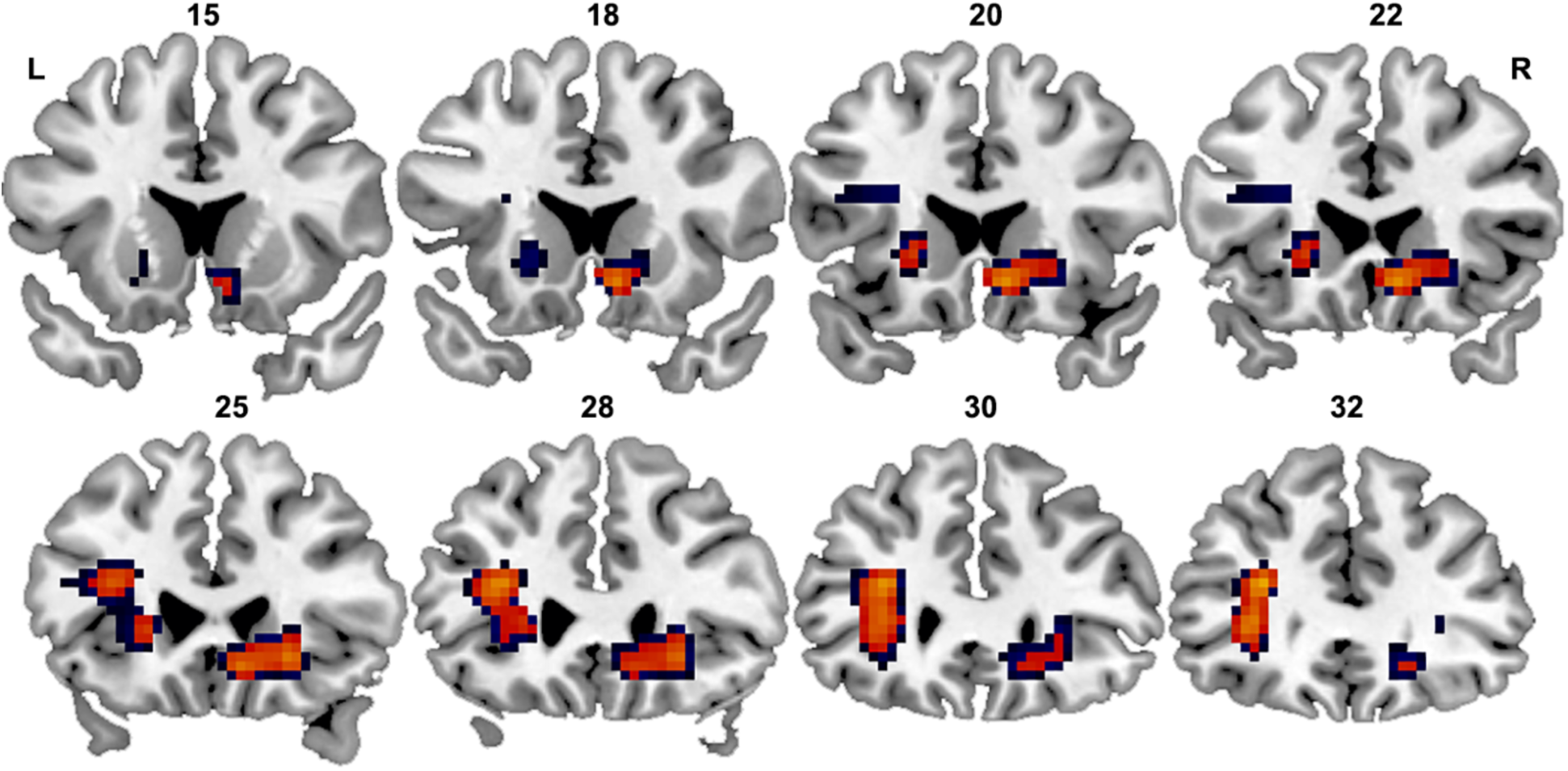
Spatial location of Iron by Age interaction on modulation of BOLD activation to parametrically increasing n-back load. Clusters exhibiting a significant (left lateralized) and trend (right lateralized) interaction between age and iron on BOLD modulation. Two height thresholds are presented here (warm scale: *p* < .001; cool scale: *p* < .005) to demonstrate the full extent of the effect for demonstration purposes. However, only a conservative height threshold of *p* = .001 was used for reporting of *FWE* cluster corrected *p*-values. Both left and right lateralized clusters extend to striatal and frontal regions. Numbers represent coronal Y-slice location along a posterior to anterior gradient; L = left, R= Right, images displayed using neurological convention.

## 3. Results

### 3.1 Behavioral Data

Accuracy and response time (RT) were recorded for all trials. To examine performance differences across WM load levels and potential age effects, two repeated-measure general linear models (GLM) were tested with WM load serving as a 4-level within-subject repeated measure and age (mean-centered) as a continuous between-subjects measure predicting either mean accuracy or median RT. For accuracy, we found significant effects of age, *F* (1,164) = 57.97, *p* < .001, and WM load, *F* (3,492) = 289.01, *p* < .001, such that accuracy declined with both increasing age and increasing WM load. A significant age × WM load interaction, *F* (3,492) = 13.63, *p* < .001, indicated that, while there were significant declines in performance across age for all levels of the *n*-back task (*p*s < .05), there were steeper declines across age for 2-, 3-, and 4-back as compared to 0-back (*p*s < .001). Response time increased with increasing WM load, *F* (3,492) = 333.81, *p* <.001, as well as with age, *F* (1,164) = 9.27, *p* =.003. There was a marginally significant age x WM load interaction, *F* (3,492) = 2.57, *p* =.054, indicating that the age effect tended to be stronger for 2-, *t*(492) = 2.40, *p* = .017, and 3-back, *t*(492) = 1.82, *p* = .069, as compared to 0-back, but similar for 4-back, *t*(492) = 0.43, *p* = .670, as compared to 0-back. Note that all trials were included (i.e., match and non-match).

### 3.2 Overall Effect of Task on BOLD Modulation

We first sought to establish the brain regions that activate and deactivate in response to parametrically increasing WM load by testing overall positive and negative modulation effect in the context of the full model with all covariates. The positive modulation effect represents regions that increased in modulation with increasing WM load, and included bilateral anterior cingulate gyrus, dorsolateral prefrontal cortex, middle frontal gyrus, superior precuneus, caudate, cerebellum and posterior parietal cortex, which corresponds well with the canonical cognitive control and working memory networks (Cabeza and Nyberg 2000). The negative modulation effect represents regions which increased in negative modulation (i.e., increased deactivation to difficulty) with increasing WM load and includes posterior cingulate gyrus, medial prefrontal cortex, superior temporal gyrus, and lateral occipital cortex; regions overlapping with task-negative and default networks (Greicius et al. 2003). This pattern of activation replicates our previous finding (Kennedy et al. 2017) in a largely overlapping sample.

### 3.3 Effect of Iron and Age on BOLD Modulation to WM Load

Contrasts of interest (effect of iron, age, and iron × age interaction) were computed from the full model that included covariates of no interest: sex, CBF, and task response time (note that the pattern of result significance is unchanged in a model excluding these covariates). There was an effect of age on BOLD modulation in response to increasing WM load in a single cluster in the fusiform gyrus (MNI xyz = 24, −39, −12, *k* = 119, *p* = .044). The effect indicated that increasing age was associated with decreased negative modulation. There were no significant main effects of iron on BOLD modulation in either the positive or the negative direction. However, as hypothesized there was a significant age × iron interaction on BOLD modulation (*p* = .001, FWE *p* < .05) in a cluster centered primarily in left putamen, caudate, and inferior frontal gyrus, and a trend for the same interaction in the right lateralized homologous regions (see **Figure 2** and **Table 2**). We present in Figure 2 results derived from a relaxed height threshold of *p* = .005, to assess whether these clusters were isolated to the fronto-striatal regions. Interestingly, no additional brain regions were present; suggesting that this interaction between age and striatal iron on brain modulation is isolated to basal ganglia and inferior frontal regions.

**Table 2.**
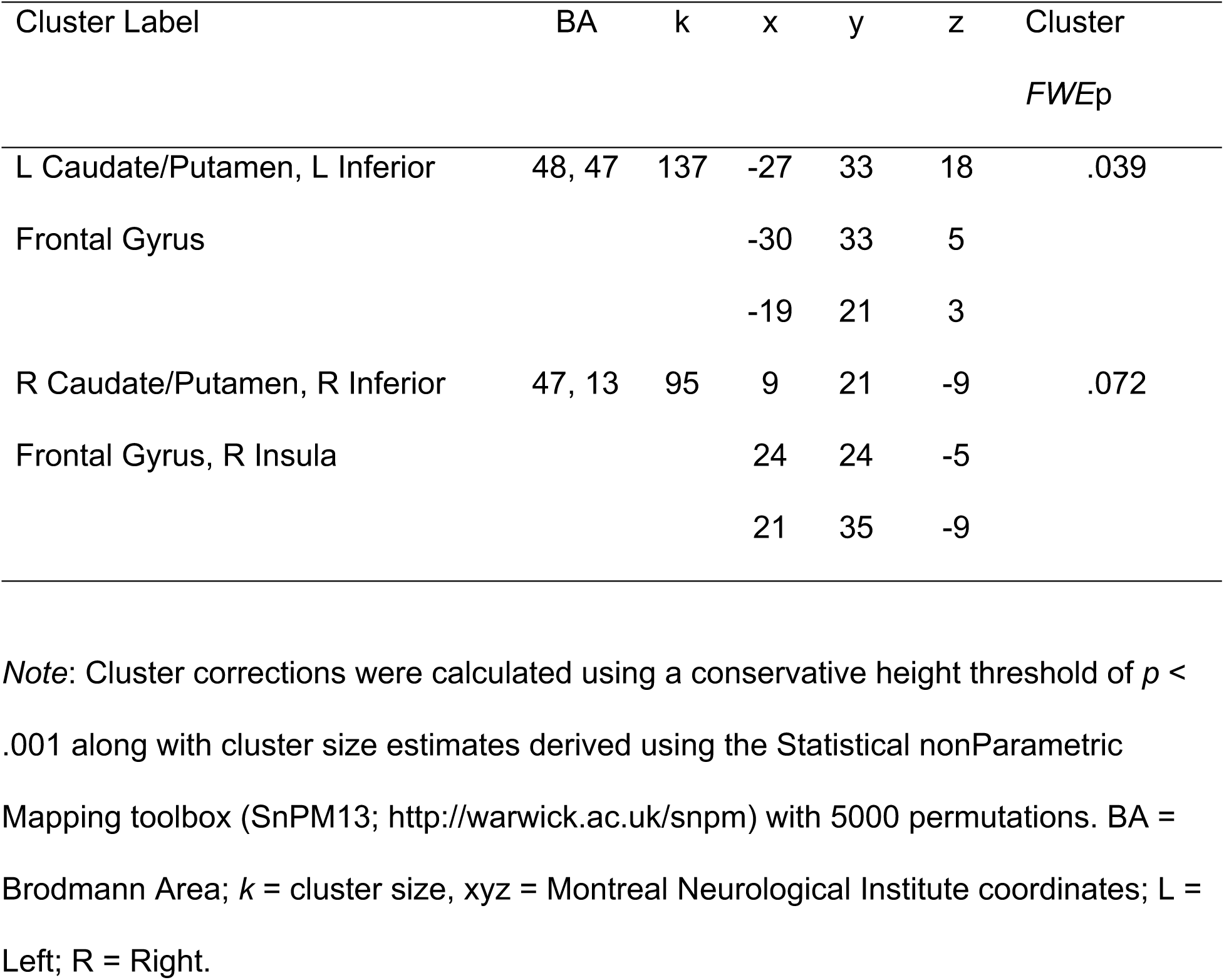
Interaction between age and iron on modulation of BOLD activation to increasing N-back load.

| Cluster Label | BA | k | x | y | z | Cluster<br>*FWE<sub>p</sub>* |
| --- | --- | --- | --- | --- | --- | --- |
| L Caudate/Putamen, L Inferior | 48, 47 | 137 | -27 | 33 | 18 | .039 |
| Frontal Gyrus |  |  | -30 | 33 | 5 |  |
|  |  |  | -19 | 21 | 3 |  |
| R Caudate/Putamen, R Inferior | 47, 13 | 95 | 9 | 21 | -9 | .072 |
| Frontal Gyrus, R Insula |  |  | 24 | 24 | -5 |  |
|  |  |  | 21 | 35 | -9 |  |

*Note:* Cluster corrections were calculated using a conservative height threshold of $p < .001$ along with cluster size estimates derived using the Statistical nonParametric Mapping toolbox (SnPM13; http://warwick.ac.uk/snpm) with 5000 permutations. BA = Brodmann Area; $k$ = cluster size, xyz = Montreal Neurological Institute coordinates; L = Left; R = Right.

To probe the interaction between age and iron on BOLD modulation we extracted beta estimates from each individual from the significant left lateralized cluster. A simple slopes analysis was conducted using the extracted beta values (Preacher et al. 2006). The results indicate that greater iron content was associated with reduced BOLD modulation to WM load, and that this effect weakened as age increased (see **Figure 3**). Specifically, for adults in the mid-50s and below, this effect was statistically significant. However, after the mid-50s this relationship was nonsignificant, suggesting that functional activation in the striatal and frontal regions in middle-age and earlier is particularly sensitive to iron in the neostriatum.

**Figure 3.**
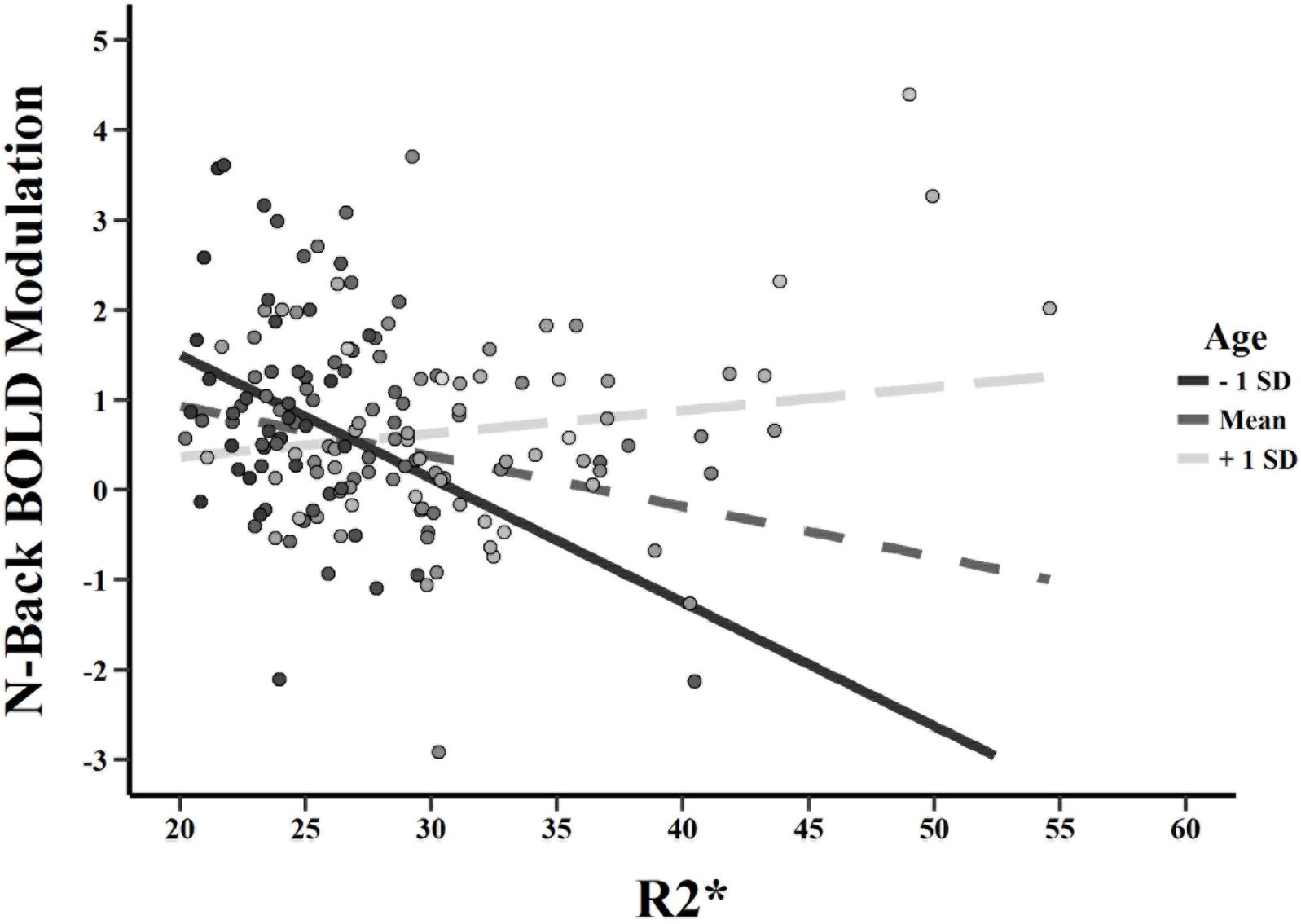
Simple slopes plot demonstrating iron (R2*) and age interaction on BOLD modulation. Beta values were extracted from the cluster exhibiting the significant interaction and simple slopes were calculated to breakdown the interaction. The effect of iron (R2*) on *n*-back modulation is plotted for three representative ages: 1 SD below the mean age (∼34 years old), the mean age (∼53 years old), and 1 SD above the mean age (∼72 years old), or young, middle-aged, and old. Higher iron content is associated with a significantly negative effect on BOLD modulation in adults below their mid-50s, however, this effect weakens as a function of older age. Shading of the dots represent the age of individual participants where the lighter colors represent increasing age. Note: SD = standard deviation.

### 3.4 Effect of Iron and BOLD Modulation on Cognition

To examine whether iron-related BOLD modulation was associated with cognitive performance we tested three models: one with task performance, one with an executive function composite, and one with a working memory composite. Age, beta estimates of modulation from the significant cluster (*p* = .001 height thresholded) and R2*served as continuous predictors (all mean-centered), along with all two-and three-way interactions as predictors of either task performance, EF, or WM. The model predicting mean task accuracy showed no iron-related BOLD modulation of task performance, *p* > .05. The working memory model showed only a trend for an interaction between age and iron on WM, *t*(158) = 1.748, β = .003, *p* = .082, but no associations between BOLD and iron on performance were found.

In contrast, the analysis with EF revealed a significant negative effect of age on EF, *t*(158) = −6.00, β = −.541, *p* < .001, and a significant positive effect of BOLD modulation on EF, *t*(158) = 3.40, β = .271, *p* < .001. These effects were qualified, however, by a marginally significant three-way interaction between modulation, iron, and age predicting EF, *t*(158) −1.94, β = −.195, *p* = .056. No other effects were significant. Simple slopes analysis was utilized to evaluate the nature of the three-way interaction (see **Figure 4**). Overall, there was a positive relationship between BOLD modulation and EF such that greater modulation to working memory load was associated with better performance on the EF composite. The interaction suggests that increasing age is associated with a weakening of the positive relationship between modulation and EF, and further, that this effect is stronger for individuals with higher iron content. Similarly, older adults with lower iron appeared to retain a strong positive relationship between modulation and EF, suggesting striatal iron may be a key modifier of the breakdown in the relationship between modulation and EF across the lifespan.

**Figure 4.**
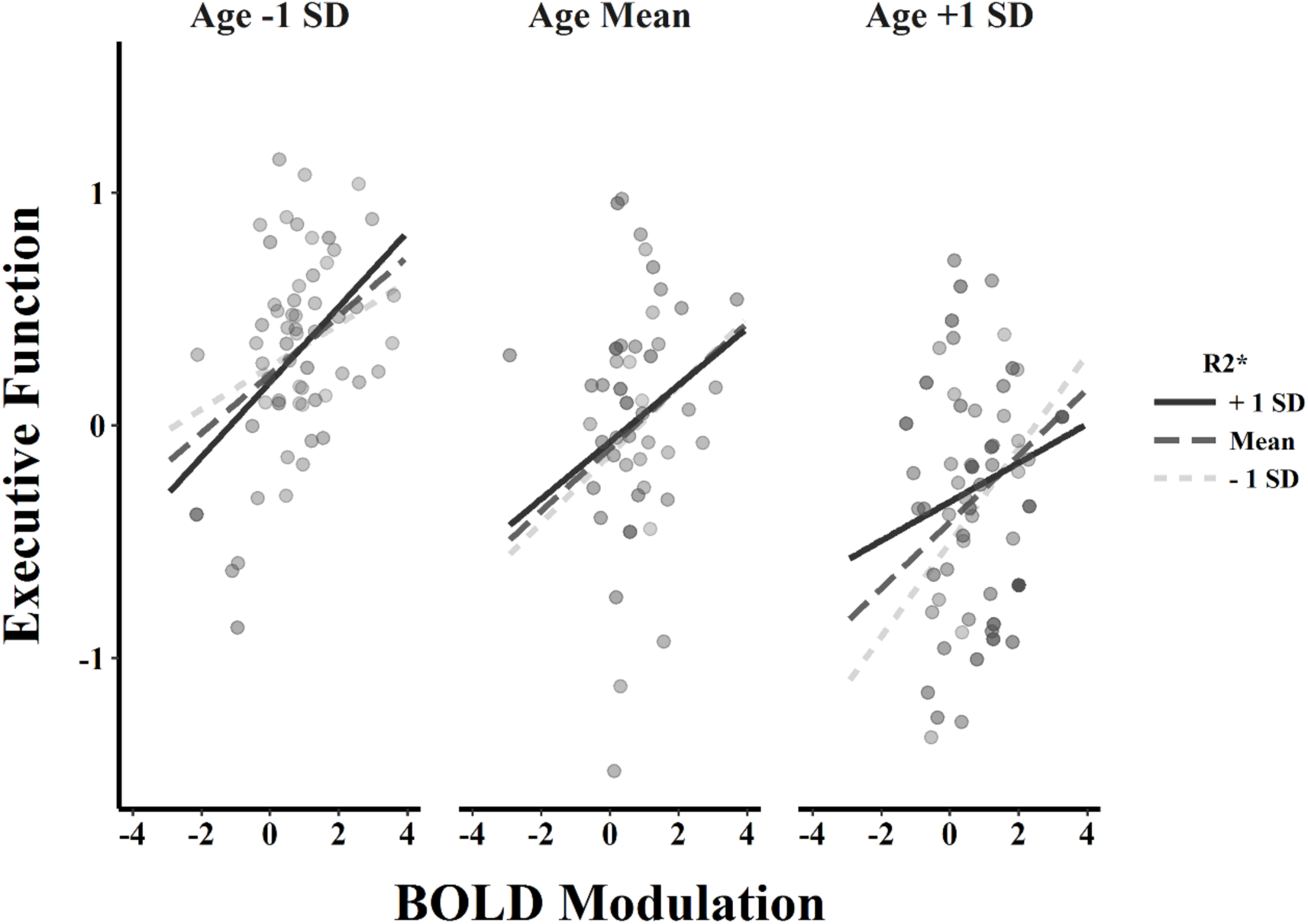
Effect of Age, Iron, and BOLD modulation on cognitive performance. A three-way interaction among R2*, BOLD modulation, and age on executive function (indexed as *z*-score composite) indicates that in general, individuals who evidence greater BOLD modulation show better executive function performance. However, this interaction suggests that increasing age is associated with a weakening of the positive relationship between modulation and EF, and further, that this effect is stronger for individuals with higher iron content. Similarly, older adults with lower iron appeared to retain a strong positive relationship between modulation and EF, suggesting striatal iron may be a key modifier of the breakdown in the relationship between modulation and EF across the lifespan. Note: SD = standard deviation.

## 4. Discussion

The present study tested the hypothesis that non-heme iron content in the striatum alters the dynamic range of age-related functional activation to increasing cognitive difficulty in a four-level *n*-back fMRI task of working memory load. We demonstrated an effect of iron concentration on BOLD response to increasing cognitive challenge, in a large contiguous cluster spanning the putamen, caudate nucleus and a portion of inferior frontal gyrus. Prominent age-related iron accumulation in astrocytes (Connor et al. 1990; Schipper 2012; Ward et al. 2014) diminishes neural signaling (Codazzi et al. 2015), which could underlie alterations in brain function as measured via fMRI. Further, iron related alterations in BOLD modulation predicted poorer cognitive performance on an executive function composite construct, suggesting that this iron-related alteration of BOLD response exerts a meaningful effect on cognition. Yet, the effect was differential by age: in middle-age and younger adults (20-50 years), age and greater iron content predicted decreased modulation of brain activation to difficulty, whereas after age 50 years (approximately) this association diminished.

These results support the findings from the first report of an effect of higher striatal iron content on task-related BOLD activity in a small group comparison of healthy young and older adults (Kalpouzos et al. 2017), where greater iron concentration was linked to decreased BOLD activation in left inferior frontal cortex and right putamen during a non-motor mental imagery task. Here we report an association between greater striatal iron concentration and modulation of BOLD signal to working memory demand in the caudate nucleus, putamen and a portion of inferior frontal gyrus. While the directionality of our finding is in accord with the previous study, differential age effects were revealed in our large, continuous lifespan sample. The differential effect of brain iron concentration on BOLD signal cannot be attributed to age-related differences in cerebral blood flow, as this was controlled statistically by an independent measurement. Rather, we find evidence in support of iron dysregulation as an early event in the neurodegenerative cascade (Hare et al. 2013).

This finding has important implications for the sensitivity of MRI estimates of brain iron concentration for early detection of cognitive deficit. Brain iron accumulation is observed continuously across the adult lifespan (Hallgren and Sourander 1958; Rodrigue et al. 2011; Daugherty and Raz 2013, 2017). However, the majority of extant evidence for its use as a biomarker included outcomes that show meaningful age-related decline after the fifth decade—namely, regional brain volume and cognitive deficit (e.g., Sullivan et al., 2009; Rodrigue et al., 2013; Daugherty et al., 2015; Daugherty & Raz, 2017). Nonetheless, prominent theories of brain iron accumulation as a harbinger of neurodegeneration, including the Free Radical Induced Energetic and Neural Decline in Senescence model (FRIENDS; Raz & Daugherty, 2018), posit functional disruption that may precede structural and cognitive decline. The current results lend credence to this notion—greater brain iron content in younger and middle-aged adults correlated with altered task-based BOLD signal and lower cognitive performance. Thus, there may be a time window, prior to the development of later-life structural decline, when brain function is sensitive to the effects of iron accumulation.

The role of non-heme iron in brain function is complex and evidences a paradoxical effect on cell function and metabolism. Under normal conditions, non-heme iron is required for ATP generation (Mills et al. 2010), dopamine synthesis and receptor expression (Youdim and Yehuda 2000), and neuronal signal transduction depends upon iron- and cytokine-related free-radicals (Lander 1997). Further, myelination processes that ensure the integrity of functional connections between the neostriatum and frontal cortex require large iron stores, presumably to meet metabolic demand (Todorich et al. 2009). However, excessive accumulation of non-heme iron outside of binding complexes will produce oxidative stress and promote chronic inflammation, which in turn impair mitochondrial energetics (Mills et al. 2010), disrupt dopamine metabolism (Meiser et al. 2013) and contribute to decline in myelin integrity (Bartzokis 2011). Therefore, non-heme iron is required for normal function but its excessive accumulation is expected to disrupt cellular and metabolic functions that, jointly, can be manifested in differences in fMRI BOLD activation, and further validated by association to lower cognitive task performance.

MRI-derived quantification of iron has yielded considerable evidence for its use as a biomarker that is sensitive to neural cognitive decline in healthy aging and neurodegenerative disease (Schenck and Zimmerman 2004; Ward et al. 2014). Within adult lifespan samples of healthy aging, greater brain iron concentration predicts age-related volumetric shrinkage over 2 years (Daugherty et al. 2015) and up to 7 years later (Daugherty and Raz 2016). Age-related increase in iron is detrimental to region-specific functions, including episodic memory (Rodrigue et al. 2013), working memory and executive function (Pujol et al. 1992; Bartzokis et al. 2011; Daugherty et al. 2015; Ghadery et al. 2015), spatial navigation ability (Daugherty and Raz 2017), and psychomotor speed (Sullivan et al. 2009) and control (Adamo et al. 2014). Further, iron accumulation over short periods predicts age-related neurodegenerative disease progression and symptom severity (Ulla et al. 2013; Walsh et al. 2013). The current report extends this literature on healthy aging further to identify a relation of regional iron concentration with altered fMRI BOLD modulation and lower executive function performance even in middle-aged and younger adults.

These findings should be interpreted in the context of the study strengths and limitations. Strengths include first, the large sample size, evenly distributed across the adult decades, allowed hypothesis testing of differential effects as a function of chronological age, which is not testable in extreme age group studies. Second, the parametric modulation of difficulty on the functional task provides a specific measure of modulation of the frontal-striatal circuitry. Third, the manual masking procedure utilized in this study allowed exclusion of partial voluming with adjacent white matter and cerebrospinal fluid when assessing iron content. Lastly, we controlled for individual differences in perfusion, as R2* is also sensitive to heme iron concentration in circulating blood, which could be a potential confound to result interpretation otherwise. Limitations include a cross-sectional study design, which preclude tests of age-related change over time or its mediation (Maxwell and Cole 2007). Second, R2* estimates are not specific measures of non-heme iron concentration because myelinated fibers can also increase R2* (Haacke et al. 2005; Glasser and Van Essen 2011). However, R2* is a well-validated index of brain iron concentration (Thomas et al. 1993; Vymazal et al. 1995), including in the striatum where few myelinated fibers are present.

In sum, striatal iron content dampens the dynamic range of functional BOLD modulation to increasing working memory load in a focal and regionally-specific manner. Through middle-age, greater striatal iron content was associated with lesser modulation in the striatum and inferior frontal gyrus in response to difficult task demands. Intriguingly, this effect was diminished in older adults, approximately after age 50 years. The linear diminishment of the effect of brain iron concertation on BOLD modulation as a function of chronological age speaks to myriad time-dependent and interactive processes that are hypothesized to drive both neural and cognitive decline. Together, these results link iron homeostasis and metabolism to brain function and cognitive outcomes, suggesting that iron content may be a sensitive biomarker of altered brain function in mid-life, and a correlate of cognitive function throughout the lifespan. Future, longitudinal research is required to understand if brain iron accumulation precipitates later, significant decline in brain function.

## Acknowledgements

This study was funded, in part, by grants from the National Institutes of Health R00 AG-036848, R00 AG-036818, R01 AG-056535, R01 AG-057537, Alzheimer’s Association NIRG-397220, as well as support from BvB Dallas and AWARE. We thank Chen Gonen for assistance with the ASL data processing and Colleen McNamee for assistance with the R2* measurements.

